# NFκB dynamics determine the stimulus-specificity of epigenomic reprogramming in macrophages

**DOI:** 10.1101/2020.02.18.954602

**Authors:** Quen J. Cheng, Sho Ohta, Katherine M. Sheu, Roberto Spreafico, Adewunmi Adelaja, Brooks Taylor, Alexander Hoffmann

**Affiliations:** Department of Microbiology, Immunology, and Molecular Genetics, University of California, Los Angeles, CA 90095; Department of Medicine, Division of Infectious Diseases, David Geffen School of Medicine, University of California, Los Angeles, CA 90095; Institute for Quantitative and Computational Biosciences, University of California, Los Angeles, CA 90095

## Abstract

The epigenome defines the cell type, but also shows plasticity that enables cells to tune their gene expression potential to the context of extracellular cues. This is evident in immune sentinel cells such as macrophages, which can respond to pathogens and cytokines with phenotypic shifts that are driven by epigenomic reprogramming^1^. Recent studies indicate that this reprogramming arises from the activity of transcription factors such as nuclear factor kappa-light-chain-enhancer of activated B cells (NFκB), which binds not only to available enhancers but may produce *de novo* enhancers in previously silent areas of the genome^2^. Here, we show that NFκB reprograms the macrophage epigenome in a stimulus-specific manner, in response only to a subset of pathogen-derived stimuli. The basis for these surprising differences lies in the stimulus-specific temporal dynamics of NFκB activity. Testing predictions of a mathematical model of nucleosome interactions, we demonstrate through live cell imaging and genetic perturbations that NFκB promotes open chromatin and formation of *de novo* enhancers most strongly when its activity is non-oscillatory. These *de novo* enhancers result in the activation of additional response genes. Our study demonstrates that the temporal dynamics of NFκB activity, which encode ligand identity^3^, can be decoded by the epigenome through *de novo* enhancer formation. We propose a mechanistic paradigm in which the temporal dynamics of transcription factors are a key determinant of their capacity to control epigenomic reprogramming, thus enabling the formation of stimulus-specific memory in innate immune sentinel cells.

## Body Text

The cellular epigenome, a regulatory network involving chromatin architecture and histone modifications, contains stable, heritable information that determines cell type-specific programs of gene expression^4^. At the same time, the epigenome of differentiated cells remains highly plastic^5,6^, particularly in immune cells like macrophages. These immune sentinel cells detect and “remember” environmental signals through epigenomic reprogramming in order to coordinate immune responses that are both context and stimulus-appropriate^1^. At a molecular level, this reprogramming is initiated by the activity of signal-dependent transcription factors (TFs) such as NFκB^7^. In cooperation with pioneer factors such as Pu.1, signal-dependent TFs increase chromatin accessibility and positive regulatory histone marks at previously latent enhancers, thus forming *de novo* enhancers^2,8^. NFκB activated by LPS has been the best-studied TF in this field. However, the degree to which NFκB or other TFs can alter the epigenome in response to different stimuli is not known.

To investigate the stimulus-specificity of *de novo* enhancer formation, we stimulated bone marrow-derived macrophages (BMDMs) with five well-characterized ligands: TNF (signaling through TNFR), Pam3CSK (TLR1/2), CpG (TLR9), LPS (TLR4), and Poly(I:C) (TLR3). We performed H3K4me1 ChIP-seq to define stimulus-dependent *de novo* enhancers and identified 3978 regions of the genome that segregated into two clusters. (Fig. 1a). The enhancers in Cluster 1 were most strongly induced by LPS and Poly(I:C) and were enriched for IRF and ISRE motifs (Fig. 1a), consistent with the fact that these stimuli activate IRF3 and type I interferon via the signaling adaptor TRIF^9^. In *Irf3*^*-/-*^*Ifnar*^*-/-*^ BMDMs these regions no longer acquired H3K4 methylation in response to LPS and Poly(I:C) (Fig. 1c). Weak induction in response to TNF was consistent with the observation that TNF does not induce IRF3 but IRF1^10^.

**Figure 1:**
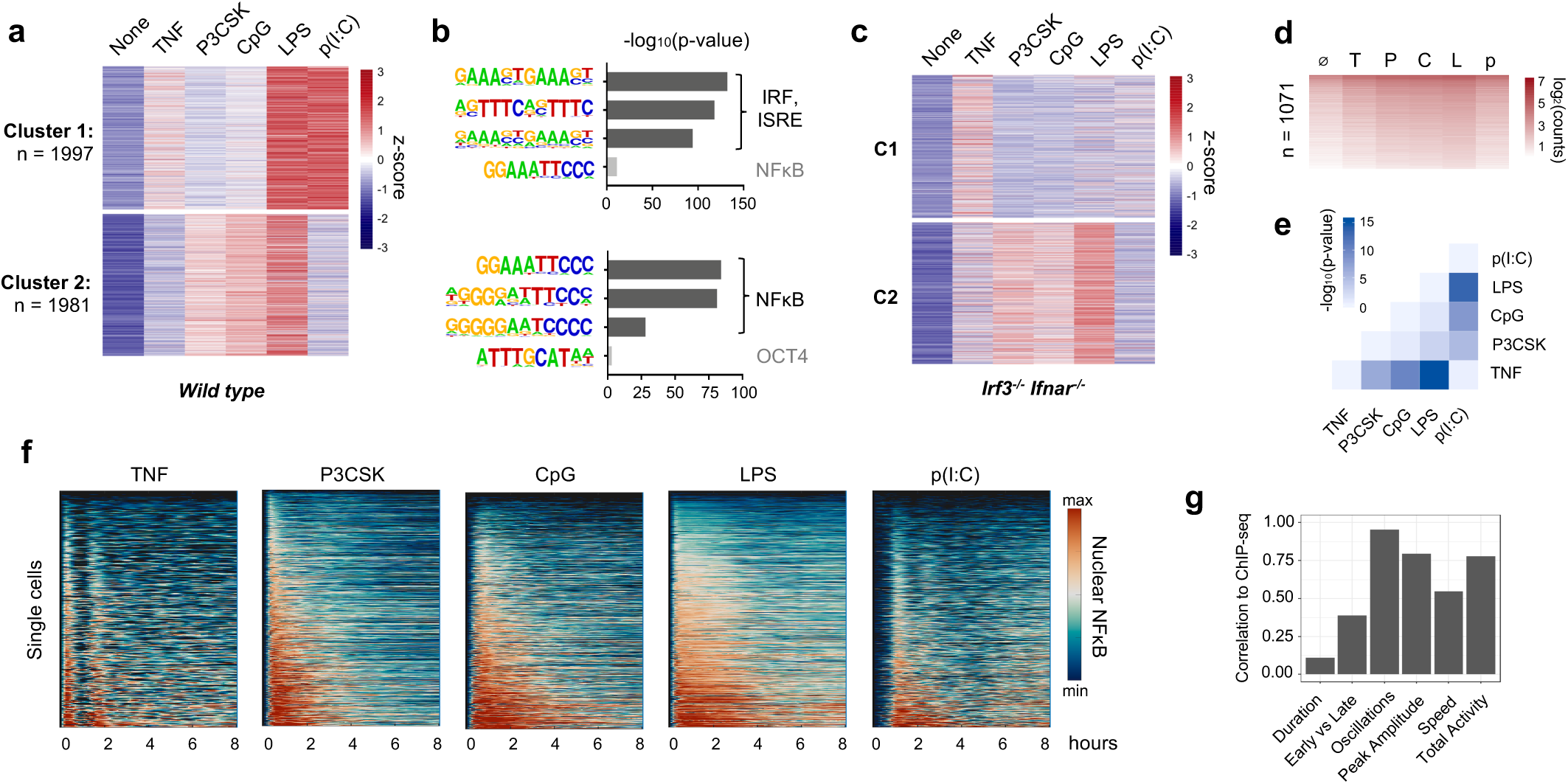
NFκB-driven de novo enhancers are stimulus-specific and correlate to dynamic features of NFκB activity. **a)** Heat map of H3K4me1 ChIP-seq inducible peaks from BMDMs stimulated with five ligands for eight hours, unsupervised K-means clustering. **b)** Known transcription factor motifs with greatest enrichment in Cluster 1 and Cluster 2 peaks. **c)** Heat map of H3K4me1 ChIP-seq in *Irf3*^-/-^*Ifnar*^-/-^ BMDMs, using same clusters as panel (a). **d)** Heat map of subset of Cluster 2 peaks that overlap with a RelA binding event by ChIP-seq. **e)** Heat map of matrix of *p*-values between ChIP-seq counts in panel (d), by two-tailed t-test between pairs of conditions. **f)** Heat maps of NFκB activity in single cells by live cell microscopy of *mVenus-RelA* BMDMs, showing nuclear abundance of NFκB in response to five stimuli. **g)** Bar graph of correlations (absolute value) between mean ChIP-seq counts in panel (d) and the six key features of NFκB dynamics^3^ (see also Extended Data, Fig. 2).

In contrast, the enhancers in Cluster 2 were highly enriched for NFκB motifs. Surprisingly, although all five stimuli activate NFκB^11^, these regions seemed to acquire H3K4me1 in a stimulus-specific manner. We observed that TNF and Poly(I:C) had little effect on these regions, while Pam3CSK, CpG, and LPS produced prominent gains in H3K4me1. These differences were consistent across replicates (Extended Data Fig. 1) and were preserved in *Irf3*^*-/-*^*Ifnar*^*-/-*^ BMDMs (Fig. 1c). Furthermore, 1071 of these regions contained an NFκB-RelA ChIP-seq peak^12^ (Fig. 1d). We concluded that these 1071 *de novo* enhancers were highly likely to be NFκB-driven. A pairwise comparison between samples quantitatively confirmed the stimulus-specificity of these enhancers (Fig. 1e), as the ChIP-seq signals of Pam3CSK, CpG, and LPS were significantly different from TNF or Poly(I:C) (*p* < 10^−5^) in these regions.

These differences would be difficult to explain if NFκB were a binary on-off switch, but NFκB is in fact activated with complex, stimulus-specific temporal dynamics^11,13,14^. Using live-cell microscopy of macrophages from mVenus-RelA mice^3^, we characterized the single-cell dynamics of NFκB p65 in response to all five ligands (Fig. 1f). We have previously identified six essential features of NFκB dynamics that function as “code words” to encode ligand identity and dose^3^. We correlated mean H3K4me1 counts in the NFκB-driven enhancers with these six features: duration, early vs late activity, oscillatory power, peak amplitude, activation speed, and total activity (Extended Data Fig. 2). We found that oscillatory power (r = −0.95), total activity (r = 0.77), and peak amplitude (r = 0.78) were correlated with the capacity of a given stimulus to form *de novo* enhancers (Fig. 1g).

We hypothesized that temporal patterns of NFκB activity might regulate its interaction with chromatin. Crystallographic studies imply that stable NFκB-DNA binding requires the DNA to be nucleosome-free, because NFκB dimers embrace the DNA double helix circumferentially^15,16^ (Fig. 2a). However, NFκB is capable of at least transiently interacting with nucleosomal DNA^17^, and can displace nucleosomes in cooperation with pioneer factor Pu.1^2^ or remodeling machinery such as SWI/SNF complexes^18^. Furthermore, the DNA-histone interface is not static but is composed of low-affinity interactions that promote spontaneous disassociation or “breathing”^19^. Thus, successive disruptions of DNA-histone contacts by NFκB may displace the nucleosome (Fig. 2b), and be followed by binding of lineage-determining TFs such as Pu.1 and the deposition of histone modifications marking the region as a *de novo* enhancer^2^.

**Figure 2:**
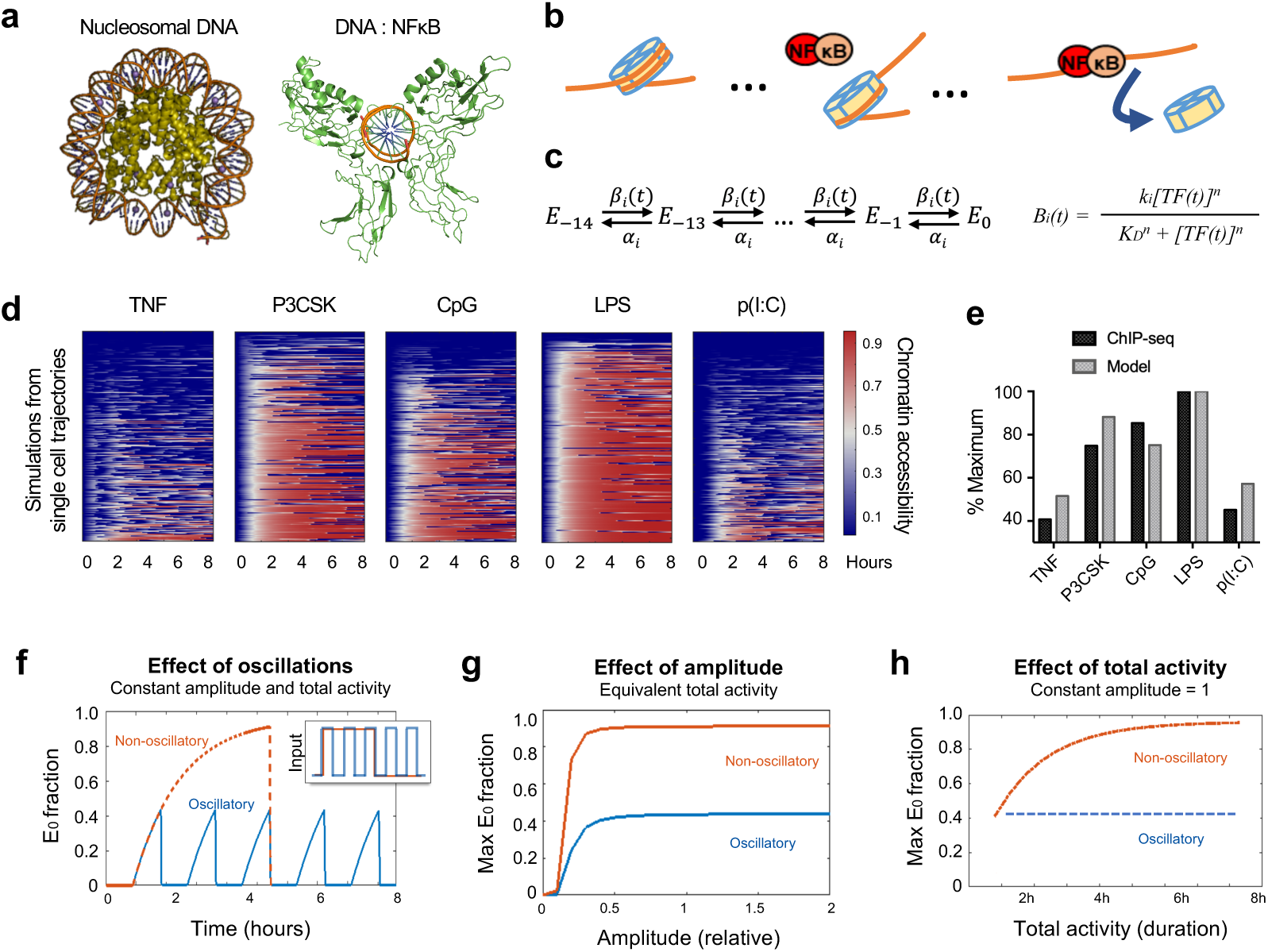
Mathematical model predicts epigenetic response to distinct dynamic features of NFκB. **a)** Crystal structures of nucleosomal DNA (PDB 1F66) *vs*. NFκB-bound DNA (PDB 1VKX), where p65:p50 NFκB dimer is in green. **b)** Schematic of model illustrating NFκB-driven displacement of nucleosome. **c)** Multi-step model with 14 steps to complete nucleosome unwrapping, each expressed as a Hill function. **d)** Heat maps of simulations of chromatin opening in response to different stimuli, using single cell trajectories from microscopy data as input. **e)** Model simulation vs. ChIP-seq data. Mean ChIP-seq counts from Fig. 1a Cluster 2, background-subtracted and scaled to maximum signal (LPS stimulation). Model simulations are mean of maximum E_0_ fraction per cell (cf. Extended Data Fig 3a), scaled to LPS condition. **f)** Model simulation of predicted chromatin accessibility comparing oscillatory *vs*. non-oscillatory input activities. **g-h)** Model simulation of predicted chromatin opening across a range of amplitudes and durations.

This provided the mechanistic basis for a multi-step model describing how dynamical NFκB activity might affect chromatin. We constructed a series of 14 Hill equations describing the competition between NFκB and histone for interacting with DNA (Fig. 2c) based on the number of contact points in the histone-DNA crystal structure^20^. Relative rates of nucleosome wrapping and unwrapping were based on available biophysical data^21^. Using measured single-cell NFκB activities (Fig. 1e) as inputs, the model simulations reproduced the differences in experimental ChIP-seq data (Fig. 2d-2e and Extended Data Fig. 3a) across a range of parameter values (Extended Data Fig. 4).

We used the model to investigate which features of NFκB dynamical activity were the key determinants of chromatin state. We examined the contribution of the three features most highly correlated with the ChIP-seq data (Fig. 1g): oscillations, amplitude, and total activity. We compared oscillatory *vs.* non-oscillatory activity while holding amplitude and total activity constant, and the model predicted that a non-oscillatory dynamic produces a two-fold greater chromatin accessibility than an oscillatory dynamic (Fig. 2f). NFκB activity must have a minimal amplitude (Fig. 2g) and extend for a minimal duration (Fig. 2h) to open chromatin. Above these thresholds, non-oscillatory NFκB has greater capacity to open chromatin than oscillatory NFκB for any given value of amplitude or duration. These simulations indicated that dynamic features of NFκB, especially the presence or absence of oscillations, determine whether it preserves or alters the chromatin state.

To test this prediction, we generated a knockout mouse in which NFκB dynamics are perturbed. In response to TNF, NFκB rapidly induces expression of *Nfkbia*, whose gene product is the negative regulator IκB*α*^22^ (Fig. 3a) and mediates oscillatory behavior of NFκB. As IκB*α* knockout mice are embryonic lethal due to chronic hyperinflammation^23^, we bred the *Nfkbia*^*-/-*^ allele into a *Rel*^*-/-*^*Tnf*^*-/-*^*Nfkbie*^*-/-*^ background, enabling the isolation of BMDMs from adult IκB*α*^-/-^ mice.

**Figure 3:**
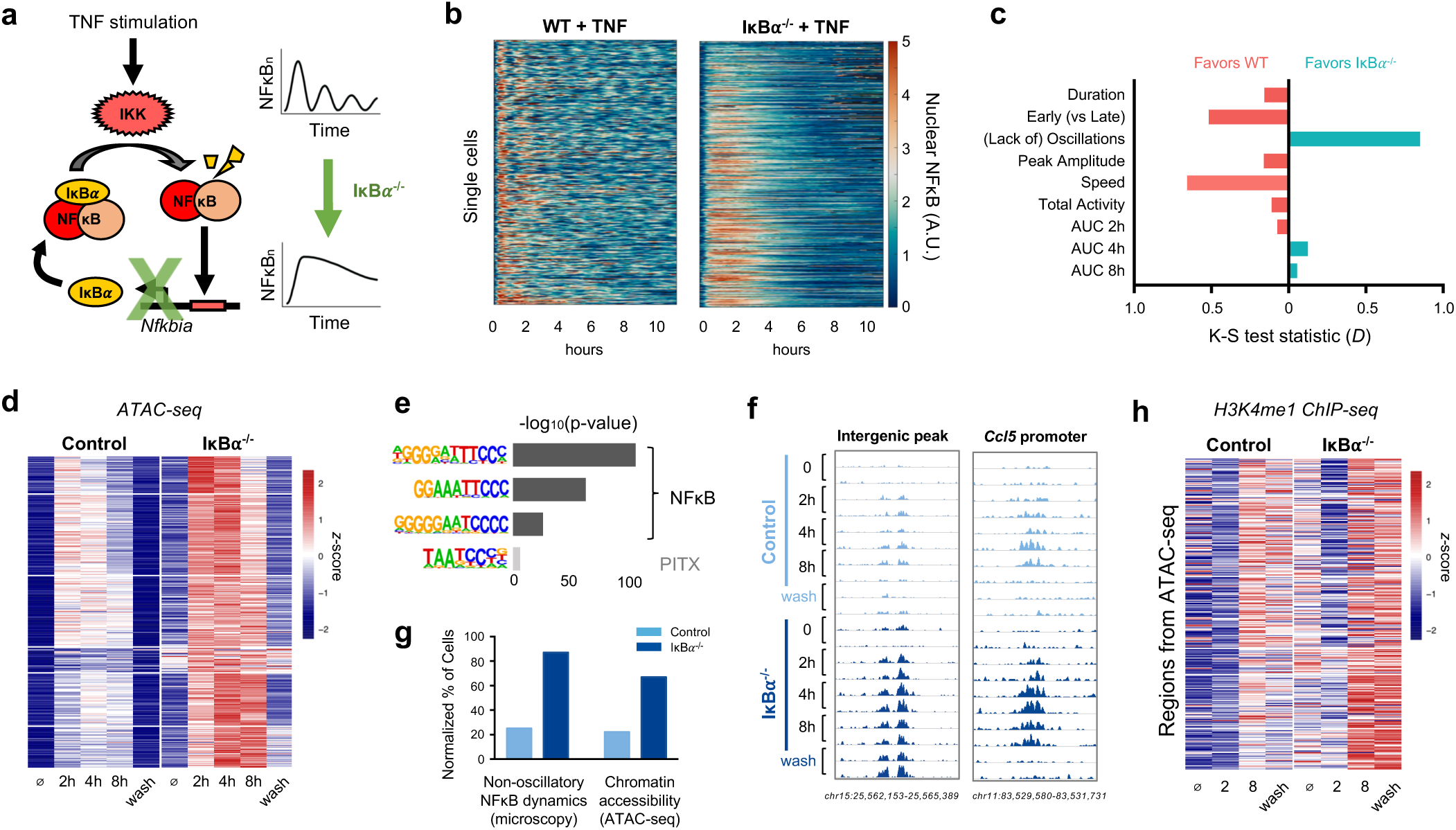
IκB*α* knockout abolishes NFκB oscillations, increasing chromatin accessibility and *de novo* enhancer formation. **a)** Schematic of IκB*α* as key regulator of NFκB oscillations. **b)** Heat map of single cell NFκB activity by microscopy comparing TNF response in WT *vs.* IκB*α*^-/-^ macrophages. **c)** Bar graph of K-S test statistic for difference in distribution of six key signaling features and areas under NFκB activity curve, comparing IκB*α*^-/-^ and WT. **d)** Heat map of ATAC-seq signal at 322 genomic regions that are TNF-inducible and differential between IκB*α*^-/-^ and control. “Wash” = 16h washout. **e)** Known transcription factor motifs with greatest enrichment in differentially inducible ATAC-seq regions. **f)** Genome browser tracks for representative differentially inducible ATAC-seq regions, two replicates per time point. **g)** Percentage of cells with non-oscillatory NFκB trajectories by microscopy, compared with relative percentage of cells with accessible chromatin at *Chr15* intergenic peak by ATAC-seq. **h)** Heat map of H3K4me1 ChIP-seq signal over the 322 regions defined as differentially inducible by ATAC-seq. “Wash” = 16h washout.

We examined the dynamics of NFκB in IκB*α*^-/-^ BMDMs by crossing these mice with *mVenus-RelA* knock-in mice and performing live cell imaging of BMDMs stimulated with TNF. We observed that knockout of IκB*α* significantly disrupted NFκB dynamics (Fig. 3b). We quantified the differences in the distribution of single cell dynamic features by Kolmogorov–Smirnov (K-S) test (Fig. 3c, Extended Data Fig. 5a) and found that the greatest dynamic difference between IκB*α*^-/-^ and WT was a loss of oscillatory activity, with a K-S test statistic (*D*) of 0.85, corresponding to a *p-*value < 10^−16^. The other key dynamic features were either unaffected, or in the case of activation speed (D = 0.66) and early-vs-late activity (D = 0.52) would intuitively favor NFκB activity in WT cells. In addition, we calculated the area under the NFκB activity curve at the time points used in subsequent experiments and found no difference (Extended Data Fig. 5b). Based on single-cell microscopy measurements, we concluded that the primary impact of IκB*α* knockout was loss of oscillations.

To profile the chromatin state, we stimulated BMDMs from IκB*α*^-/-^ and littermate controls with TNF and performed ATAC-seq at two, four, and eight hours. This was followed by a 16-hour washout period, and a final time point was collected after washout (Extended Data Fig. 6a). We identified 1443 genomic regions that demonstrated TNF-inducible chromatin accessibility in either genotype. Of these, 332 were differentially inducible between control and IκB*α*^-/-^. Strikingly, 97% of these regions (n=322) had greater chromatin accessibility in the knockout than control (Fig. 3d). These differentially inducible regions were strongly enriched for NFκB motifs (Fig. 3e), and 311 of 322 overlapped with a RelA ChIP-seq peak (Extended Data Fig. 6c). Differentially inducible regions were more likely than constitutively accessible regions to fall in intergenic portions of the genome (Extended Data Fig. 6b), suggesting that they tend to function as *cis*-acting enhancer elements near key innate immune genes such as *Ccl5* (Fig. 3f), which has previously been shown to require chromatin remodeling for full induction^12^.

Our model predicted that chromatin accessibility is primarily determined by whether NFκB is oscillatory or non-oscillatory within a single cell. We therefore considered that the magnitude of ATAC-seq signal can be interpreted as the proportion of cells in a sample in which a particular region of DNA is accessible. By microscopy, 87% of IκB*α*^-/-^ cells have non-oscillatory NFκB, compared to 25% in WT cells. This was similar to the magnitude of ATAC-seq differences between IκB*α*^-/-^ and control. For example, at an intergenic peak on chromosome 15, 67% of the cells in IκB*α*^-/-^ were accessible, compared to 22% of cells in the control (Fig. 3g).

To investigate more definitively that the negative feedback function of IκB*α* rather than its basal activity is critical for the observed effects, we utilized a recently described IκB*α*^κB/κB^ mutant in which NFκB-binding sites in the promoter of the *Nfkbia* gene are disrupted^24^ (Extended Data Fig. 7a). In this model, basal IκB*α* expression is preserved, and the mice live into adulthood without requiring compound suppressor mutations. We confirmed that upon TNF stimulation IκB*α*^κB/κB^ BMDMs activate NFκB in a non-oscillatory manner (Extended Data Fig. 7b). ATAC-seq analysis of TNF-stimulated WT vs IκB*α*^κB/κB^ BMDMs recapitulated our findings in the IκB*α*^-/-^ system, with 131 genomic regions demonstrating greater gain of chromatin accessibility in the mutant compared to WT (Extended Data Fig. 7c). These regions were enriched for NFκB motifs, and 90% overlapped with a RelA ChIP-seq peak (Extended Data Fig. 7d-7e). Taken together, the ATAC-seq data from both IκB*α*^-/-^ and IκB*α*^κB/κB^ experimental models indicated that loss of inducible negative feedback in the NFκB signaling system, which results in a loss of oscillations, results in greater chromatin accessibility.

Next, we examined whether regions with differentially inducible chromatin accessibility acquire the corresponding histone mark of enhancers. We performed H3K4me1 ChIP-seq in TNF-stimulated control and IκB*α*^-/-^ BMDMs and found that in the 322 differentially inducible ATAC-seq regions there was also a greater gain of H3K4me1 signal in IκB*α*^-/-^ than control (Fig. 3h). Notably, these histone marks persisted even after a 16-hour washout. This suggests that chromatin opening facilitated by NFκB may be transient but leads to durable H3K4 methylation even after the stimulus is removed, marking the region as a *de novo* enhancer and reprogramming the epigenome.

Because histone methylation is more durable and indicative of enhancer function, we analyzed the H3K4me1 ChIP-seq data independently and identified 2081 regions that acquired more H3K4 methylation in IκB*α*^-/-^ than control (Fig. 4a). These differentially induced, dynamics-dependent *de novo* enhancers persisted after the TNF stimulus was washed out, and they were strongly enriched for NFκB motifs (Fig. 4b). We then asked whether these regions, which are dependent on non-oscillatory NFκB in the IκB*α*^-/-^ system, corresponded to the stimulus-specific NFκB-driven *de novo* enhancers in WT BMDMs (Fig. 1d). We found that there was a highly significant overlap (*p* = 10 e-45), and the inducible ChIP-seq signal was consistently greater when NFκB dynamics were non-oscillatory rather than oscillatory, whether by genetic perturbation or by stimulus-specific signaling mechanisms (Fig. 4c).

**Figure 4:**
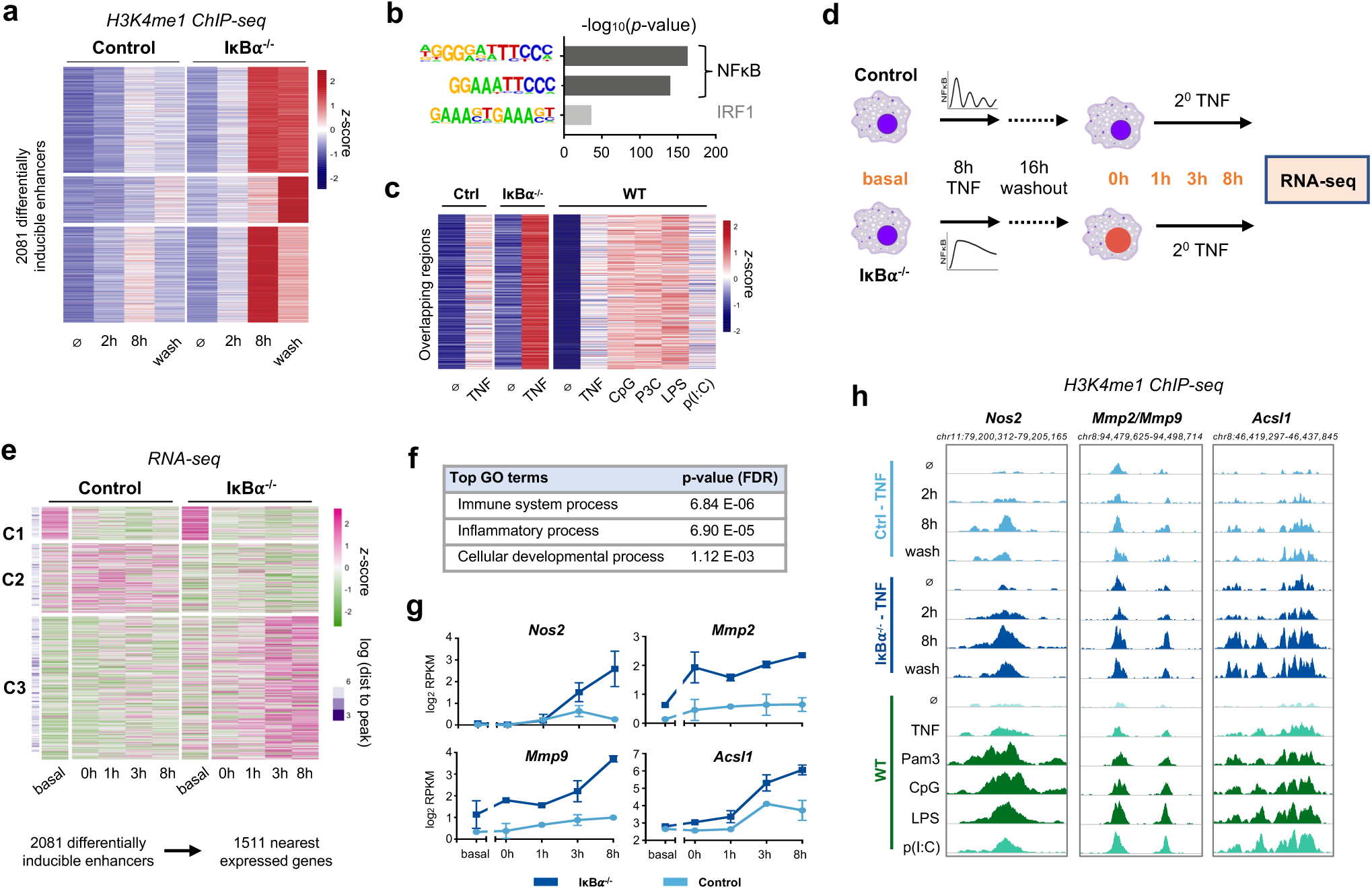
NFκB dynamics-dependent enhancers are associated with dynamics-dependent gene expression. **a)** Heat map of H3K4me1 ChIP-seq signal at 2081 regions that are TNF-inducible and differential between IκB*α*^-/-^ and control, i.e. dynamics-dependent enhancers. “Wash” = 16h washout. **b)** Known transcription factor motifs with greatest enrichment in dynamics-dependent enhancers. **c)** Heat map of H3K4me1 signal after 8h stimulation at regions that overlap between Fig. 4a and Fig. 1d (n=211, *p* for overlap=10 e-45). **d)** Schematic of RNA-seq experiment. **e)** Heat map showing expression of genes closest to dynamics-dependent enhancers, where Cluster 3 exhibits differential gene expression between IκB*α*^-/-^ and control. **f)** Top biological process ontology terms for genes in Cluster 3 of Fig. 4e. **g)** Examples of genes differentially induced between IκB*α*^-/-^ and control, average and standard deviation of two replicates. **h)** Genome browser tracks of differentially inducible H3K4me1 peaks near differentially inducible genes, showing TNF-stimulated IκB*α*^-/-^ *vs.* control and stimulus-specific response in WT BMDMs. More darkly shaded tracks indicate non-oscillatory NFκB conditions.

Next, we asked whether these NFκB dynamics-dependent enhancers had a functional role in macrophage gene expression. We hypothesized that *de novo* enhancers would alter transcriptional responses to subsequent stimulation. We primed control and IκB*α*^-/-^ BMDMs with TNF for eight hours followed by 16-hour washout as before, then re-stimulated with secondary TNF over eight hours (Fig. 4d). We performed RNA-seq in the basal (untreated) condition and at zero, one, three, and eight hours of secondary TNF stimulation. We explored the relationship between differentially inducible enhancers and gene expression using two approaches. First, using a peak-centric approach, we linked the 2081 enhancers to their nearest expressed genes, removed duplicates, and identified three distinct patterns of expression for the 1511 genes. Cluster 1 and 2 genes were not TNF-responsive in either condition, reflecting an intrinsic limitation of this approach when enhancers often regulate much more distant genes^25^. Despite this limitation, 58% of nearest genes were both TNF-responsive and more strongly induced in IκB*α*^-/-^ BMDMs (Fig. 4e Cluster 3). Many of these genes were not induced in controls at all. The differentially induced genes were enriched for ontology terms “Immune system process” and “Inflammatory process” (Fig. 4f).

To corroborate the results from the peak-centric analysis, we also examined our data using a gene-centric approach. From the RNA-seq dataset we identified 1958 TNF-inducible genes, 482 of which were differentially regulated in IκB*α*^-/-^ versus control (Extended Data Fig. 8a-8b). For each gene, we measured the genomic distance to the nearest differentially inducible H3K4me1 ChIP-seq region. We found that differentially inducible genes were significantly closer to differentially inducible enhancers (*p* = 1.13 e-9) than genes that were not differentially inducible (Extended Data Fig. 8c-8d). Thus, both analytical approaches indicated that NFκB dynamics-dependent *de novo* enhancers play a functional role in differentially regulating gene expression response to secondary TNF.

The dynamics-dependent gene expression program included *Nos2, Mmp2*, and *Mmp9*, which are well-defined markers of classical macrophage activation^26^, as well as *Acsl1*, which plays a role in the pathogenesis of atherosclerosis^27^ (Fig. 4g). Each of these genes had a nearby enhancer that gained more H3K4me1 signal in the presence of non-oscillatory NFκB, whether in the IκB*α*^-/-^ system or in WT BMDMs stimulated with different ligands (Fig. 4h). These specific examples further suggested that *de novo* enhancers formed by non-oscillatory NFκB regulate genes involved in macrophage activation.

In summary, our results indicate that NFκB dynamics, particularly whether it is oscillatory or non-oscillatory, determine its capacity to reprogram the macrophage epigenome. We show with a mathematical model how biophysical principles governing nucleosome dynamics might decode stimulus-specific NFκ B dynamical features. The role of temporal dynamics may thus complement the structure-function model in which pioneering TFs access nucleosomal DNA based on their recognition of partially exposed DNA motifs^28^. More broadly, our findings imply that stimulus-specific temporal dynamics of TF activity may result in stimulus-specific memory in macrophages. In response to some stimuli, immune sentinel cells activate oscillatory NFκB, which is sufficient for gene expression but does not produce *de novo* enhancers. In response to other stimuli, cells activate non-oscillatory NFκB, which activates a comparable gene expression program^29^ while also altering the epigenome, changing the phenotypic state of the cell and its response to subsequent stimuli. While further work will be needed to determine the physiological functions of NFκB dynamics-dependent *de novo* enhancers, our study establishes a mechanistic paradigm of TF temporal dynamics being a key determinant for driving epigenetic reprogramming.

## Acknowledgements

This work was supported by NIH grants R01-AI127864, R01-GM117134, F31AI138450, T32GM008042, and T32-AI089398, as well as the Specialty Training and Advanced Research (STAR) program of the UCLA Department of Medicine. We would like to thank Diane Lefaudeux and Kensei Kishimoto for bioinformatics advice, and Eason Lin and Ying Tang for their insights and critical reading of the manuscript. Sequencing was performed at the UCLA Broad Stem Cell Center Sequencing Core.

## Author contributions

QC, SO, AA, and BT performed the experiments. QC, SO, KS, RS, and AA analyzed the data. KS and BT developed the mathematical model. QC and AH wrote the manuscript with input from KS. All authors reviewed the manuscript. AH coordinated and funded the work.

## Bibliography

1. Murray PJ, Wynn TA. Protective and pathogenic functions of macrophage subsets. Nat Rev Immunol. 2011 Oct 14;11(11):723–37. PMCID: PMC3422549

2. Ostuni R, Piccolo V, Barozzi I, Polletti S, Termanini A, Bonifacio S, Curina A, Prosperini E, Ghisletti S, Natoli G. Latent enhancers activated by stimulation in differentiated cells. Cell. 2013 Jan 17;152(1–2):157–71. PMID: 23332752

3. Taylor B, Adelaja A, Liu Y, Luecke S, Hoffmann A. Macrophages classify immune threats using at least six codewords of the temporal NFkB code. BioRxiv. 2020;

4. Allis CD, Jenuwein T. The molecular hallmarks of epigenetic control. Nat Rev Genet. 2016 Aug;17(8):487–500.

5. Glass CK, Natoli G. Molecular control of activation and priming in macrophages. Nature Immunology. 2015;17:26–33.

6. Heinz S, Romanoski CE, Benner C, Glass CK. The selection and function of cell type-specific enhancers. Nat Rev Mol Cell Biol. 2015 Mar;16(3):144–154. PMCID: PMC4517609

7. Lawrence T. The Nuclear Factor NF-B Pathway in Inflammation. Cold Spring Harbor Perspectives in Biology. 2009 Dec 1;1(6):a001651–a001651.

8. Kaikkonen MU, Spann NJ, Heinz S, Romanoski CE, Allison KA, Stender JD, Chun HB, Tough DF, Prinjha RK, Benner C, Glass CK. Remodeling of the enhancer landscape during macrophage activation is coupled to enhancer transcription. Mol Cell. 2013 Aug 8;51(3):310–325. PMCID: PMC3779836

9. Honda K, Takaoka A, Taniguchi T. Type I Inteferon Gene Induction by the Interferon Regulatory Factor Family of Transcription Factors. Immunity. 2006 Sep;25(3):349–360.

10. Yarilina A, Park-Min K-H, Antoniv T, Hu X, Ivashkiv LB. TNF activates an IRF1-dependent autocrine loop leading to sustained expression of chemokines and STAT1-dependent type I interferon-response genes. Nat Immunol. 2008 Apr;9(4):378–387. PMID: 18345002

11. Werner SL, Barken D, Hoffmann A. Stimulus specificity of gene expression programs determined by temporal control of IKK activity. Science. 2005 Sep 16;309(5742):1857–1861. PMID: 16166517

12. Tong A-J, Liu X, Thomas BJ, Lissner MM, Baker MR, Senagolage MD, Allred AL, Barish GD, Smale ST. A Stringent Systems Approach Uncovers Gene-Specific Mechanisms Regulating Inflammation. Cell. 2016 Mar 24;165(1):165–179. PMCID: PMC4808443

13. Behar M, Hoffmann A. Understanding the temporal codes of intra-cellular signals. Curr Opin Genet Dev. 2010 Dec;20(6):684–693. PMCID: PMC2982931

14. Covert MW, Leung TH, Gaston JE, Baltimore D. Achieving stability of lipopolysaccharide-induced NF-kappaB activation. Science. 2005 Sep 16;309(5742):1854–1857. PMID: 16166516

15. Chen FE, Huang DB, Chen YQ, Ghosh G. Crystal structure of p50/p65 heterodimer of transcription factor NF-kappaB bound to DNA. Nature. 1998 Jan 22;391(6665):410–413. PMID: 9450761

16. Suto RK, Clarkson MJ, Tremethick DJ, Luger K. Crystal structure of a nucleosome core particle containing the variant histone H2A.Z. Nat Struct Biol. 2000 Dec;7(12):1121–1124. PMID: 11101893

17. Lone IN, Shukla MS, Charles Richard JL, Peshev ZY, Dimitrov S, Angelov D. Binding of NF-κB to Nucleosomes: Effect of Translational Positioning, Nucleosome Remodeling and Linker Histone H1. Schübeler D, editor. PLoS Genet. 2013 Sep 26;9(9):e1003830.

18. Kobayashi K, Hiramatsu H, Nakamura S, Kobayashi K, Haraguchi T, Iba H. Tumor suppression via inhibition of SWI/SNF complex-dependent NF-κB activation. Sci Rep. 2017 Dec;7(1):11772.

19. Li G, Levitus M, Bustamante C, Widom J. Rapid spontaneous accessibility of nucleosomal DNA. Nat Struct Mol Biol. 2005 Jan;12(1):46–53.

20. Davey CA, Sargent DF, Luger K, Maeder AW, Richmond TJ. Solvent mediated interactions in the structure of the nucleosome core particle at 1.9 a resolution. J Mol Biol. 2002 Jun 21;319(5):1097–1113. PMID: 12079350

21. Tims HS, Gurunathan K, Levitus M, Widom J. Dynamics of Nucleosome Invasion by DNA Binding Proteins. Journal of Molecular Biology. 2011 Aug;411(2):430–448.

22. Hoffmann A, Levchenko A, Scott ML, Baltimore D. The IkappaB-NF-kappaB signaling module: temporal control and selective gene activation. Science. 2002 Nov 8;298(5596):1241–1245. PMID: 12424381

23. Beg AA, Sha WC, Bronson RT, Baltimore D. Constitutive NF-kappa B activation, enhanced granulopoiesis, and neonatal lethality in I kappa B alpha-deficient mice. Genes Dev. 1995 Nov 15;9(22):2736–2746. PMID: 7590249

24. Peng B, Ling J, Lee AJ, Wang Z, Chang Z, Jin W, Kang Y, Zhang R, Shim D, Wang H, Fleming JB, Zheng H, Sun S-C, Chiao PJ. Defective feedback regulation of NF-kappaB underlies Sjogren’s syndrome in mice with mutated kappaB enhancers of the IkappaBalpha promoter. Proc Natl Acad Sci USA. 2010 Aug 24;107(34):15193–15198. PMCID: PMC2930541

25. Corces MR, Granja JM, Shams S, Louie BH, Seoane JA, Zhou W, Silva TC, Groeneveld C, Wong CK, Cho SW, Satpathy AT, Mumbach MR, Hoadley KA, Robertson AG, Sheffield NC, Felau I, Castro MAA, Berman BP, Staudt LM, Zenklusen JC, Laird PW, Curtis C, The Cancer Genome Atlas Analysis Network†, Greenleaf WJ, Chang HY. The chromatin accessibility landscape of primary human cancers. Science. 2018 Oct 26;362(6413):eaav1898.

26. Murray PJ, Allen JE, Biswas SK, Fisher EA, Gilroy DW, Goerdt S, Gordon S, Hamilton JA, Ivashkiv LB, Lawrence T, Locati M, Mantovani A, Martinez FO, Mege JL, Mosser DM, Natoli G, Saeij JP, Schultze JL, Shirey KA, Sica A, Suttles J, Udalova I, van Ginderachter JA, Vogel SN, Wynn TA. Macrophage activation and polarization: nomenclature and experimental guidelines. Immunity. 2014 Jul 17;41(1):14–20. PMCID: PMC4123412

27. Kanter JE, Kramer F, Barnhart S, Averill MM, Vivekanandan-Giri A, Vickery T, Li LO, Becker L, Yuan W, Chait A, Braun KR, Potter-Perigo S, Sanda S, Wight TN, Pennathur S, Serhan CN, Heinecke JW, Coleman RA, Bornfeldt KE. Diabetes promotes an inflammatory macrophage phenotype and atherosclerosis through acyl-CoA synthetase 1. Proc Natl Acad Sci USA. 2012 Mar 20;109(12):E715–724. PMCID: PMC3311324

28. Soufi A, Garcia MF, Jaroszewicz A, Osman N, Pellegrini M, Zaret KS. Pioneer transcription factors target partial DNA motifs on nucleosomes to initiate reprogramming. Cell. 2015 Apr 23;161(3):555–568. PMCID: PMC4409934

29. Cheng CS, Behar MS, Suryawanshi GW, Feldman KE, Spreafico R, Hoffmann A. Iterative Modeling Reveals Evidence of Sequential Transcriptional Control Mechanisms. Cell Syst. 2017 Mar 22;4(3):330–343 e5. PMCID: PMC5434763

