## Supplementary material for "NFκB dynamics determine the stimulus-specificity of epigenomic reprogramming in macrophages": Methods

**Mice**

All mouse experiments were performed in a C57Bl/6 background. The mVenus-RelA reporter mouse (*Rela*$^{V/V}$), in which an mVenus-RelA fusion is knocked in to the endogenous *Rela* locus, has been previously described[1]. As *Nfkbia*$^{-/-}$ mice are perinatal lethal due to chronic inflammation they were crossed with *Rel*$^{-/-}$*Tnf*$^{-/-}$ alleles to achieve rescue and enable BMDM isolation[2]; they were also crossed with *Nfkbie*$^{-/-}$ alleles to avoid compensation by IκBε[3]. For ATAC, ChIP, and RNA-seq experiments *Rel*$^{-/-}$*Tnf*$^{-/-}$*Nfkbie*$^{-/-}$ *Nfkbia*$^{+/-}$ heterozygotes were mated, and *Nfkbia*$^{+/+}$ *and Nfkbia*$^{-/-}$ littermates were used for control and knockout, respectively. The *Nfkbia*$^{\kappa B/\kappa B}$ mice have been previously described[4] and were a gift from Dr. Paul Chiao. For live cell microscopy, *Rel*$^{+/-}$*Tnf*$^{+/-}$*Nfkbie*$^{+/-}$*Nfkbia*$^{-/-}$*Rela*$^{V/V}$ and *Nfkbia*$^{\kappa B/\kappa B}$*Rela*$^{V/V}$ mice were compared to *Rela*$^{V/V}$ controls.

**Tissue culture**

BMDMs were generated by isolating bone marrow from the femurs and tibias of sex-matched mice between the ages of six and 12 weeks. Bone marrow cells were incubated in L929-conditioned media (RPMI + 30% L929 media, 10% FBS, 100 IU/ml penicillin, 100 $\mu$g/ml streptomycin, 2 mM L-glutamine) at 37$^0$C for seven days. Cells were stimulated with TNF-$\alpha$ (10 ng/ml, R&D Systems 410-MT), Pam3CSK4 (100 ng/ml, Invivogen tlrl-pms), CpG (1 $\mu$M, Invivogen tlrl-1668), LPS (100 ng/ml, Sigma-Aldrich L2880), or Poly(I:C) (50 $\mu$g/ml, Invivogen tlrl-picw). Ligand doses were chosen to maximize differences in NFκB signaling dynamics[1].

**Live cell microscopy and analysis**

Microscopy and analysis have been described in detail previously[1,5]. Briefly, BMDMs derived from mVenus-RelA reporter mice were plated in an 8-well ibidi SlideTek chamber, stained with 2.5 ng/mL Hoechst 33342 in PBS, then stimulated with the ligands listed. Cells were imaged at 5-minute intervals on a Zeiss Axio Observer platform with live-cell incubation, using epifluorescent excitation from a Sutter Lambda XL light source. Images were recorded on a Hamamatsu Orca Flash 2.0 CCD camera. Time-lapse images were exported for single-cell tracking and measurement in MATLAB R2016a. Cells were identified using DIC images, and nuclear/cytoplasmic compartments were defined by the Hoechst image. Nuclear NFκB levels in the mVenus channel were quantified on a per-cell basis, and normalized to image background levels. Mitotic cells, dead cells, and cells that drifted out of the field of view, were excluded from analysis. Dynamic features of NFκB activity were measured for each single cell trajectory. The complete library of features and identification of six signaling “code words” was described previously[1].

**H3K4me1 ChIP-seq libraries**

BMDMs were cross-linked with 1% formaldehyde, quenched with 125 mM glycine, frozen and stored at -80$^0$C. Cell pellets were lysed in 50 mM HEPES-KOH pH 7.6, 140 mM NaCl, 1 mM EDTA, 10% glycerol, 0.5% NP-40, 0.25% Triton X-100, and 1x protease inhibitor cocktail (Thermo Scientific 78439) with one

15-second cycle of sonication in a Bioruptor (Diagenode). Nuclei were washed in 10 mM Tris-HCl pH 8.0, 200 mM NaCl, 1 mM EDTA, 0.5 mM EGTA, and 1x protease inhibitor cocktail. Nuclei were then resuspended in 10 mM Tris-HCl pH 8.0, 100 mM NaCl, 1 mM EDTA, 0.5 mM EGTA, 0.1% Na Deoxycholate, 0.5% N-lauroylsarcosine, 0.2% SDS, and 1x protease inhibitor cocktail, and subjected to twelve 30-second cycles of sonication in a Bioruptor. The resulting chromatin fragments were diluted with 5.2 volumes of 10 mM Tris-HCl pH 8.0, 160 mM NaCl, 1 mM EDTA, 0.01% SDS, 1.2% Triton X-100, 1x protease inhibitor cocktail, and incubated for 1.5 hours with Protein-G DynaBeads (Thermo Fisher 10004D) for pre-clearing. Protein G beads were removed, a 1% aliquot of DNA was taken as input, and remaining chromatin was incubated with rabbit anti-H3K4me1 antibody (Abcam ab8895) overnight at $4^0$C at a ratio of 2 $\mu$g antibody per 6 million cells of starting material.

For immunoprecipitation, 30 $\mu$L of Protein G beads were added to antibody-chromatin complexes and incubated at $4^0$C for three hours. Supernatant was removed, and beads were serially washed with Low Salt buffer (50 mM HEPES-KOH pH 7.6, 140 mM NaCl, 1 mM EDTA, 1% Triton X-100, 0.1% Na Deoxycholate, 0.1% SDS), High Salt buffer (50 mM HEPES-KOH pH 7.6, 500 mM NaCl, 1 mM EDTA, 1% Triton X-100, 0.1% Na Deoxycholate, 0.1% SDS), LiCl buffer (20 mM Tris-HCl pH 8.0, 250 mM LiCl, 1 mM EDTA, 0.5% Na Deoxycholate, 0.5% NP-40), and TE buffer (10 mM Tris-HCl pH 8.0, 1 mM EDTA). Immunoprecipitated chromatin complexes were treated with RNAse A (Thermo Fisher 12091021) at $37^0$C for one hour, and crosslinks were reversed with 10% SDS and 0.6 mg/ml Proteinase K (New England Biolabs P81075) overnight with shaking at $65^0$C. Immunoprecipitated DNA fragments were purified with AMPure XP SPRI beads (Beckman Coulter A63881) at a 0.95 volume ratio according to manufacturer's instructions.

Libraries were prepared for sequencing using NEBNext Ultra II DNA Library Prep Kit (New England Biolabs E7645) with NEBNext Multiplex Oligos (New England Biolabs E7335 and E7500). Each library was prepared from 100 ng of starting DNA. Input samples were pooled from input DNA of the same genotype. Final libraries were checked for quality by agarose gel, quantified with Qubit (Life Technologies Q32851), and multiplexed with a maximum of 24 samples per sequencing reaction.

**ATAC-seq libraries**

BMDMs were dissociated with Accutase (Thermo Fisher Scientific), and 50,000 cells were used to prepare nuclei. Cell membranes were lysed using cold lysis buffer (10mM Tris-HCl pH7.5, 3mM MgCl2, 10mM NaCl and 0.1% IGEPAL CA-630). Nuclei were pelleted by centrifugation for 10 minutes at 500 x *g* and suspended in transposase reaction mixture (25 µl of 2X TD Buffer (Illumina), 2.5 µl of TD Enzyme 1 (Illumina), and 22.5 µl of nuclease-free water). The transposase reaction was performed for 30 minutes at $37^0$C in a thermomixer shaker. Fragmented DNA in the reaction was purified using MinElute PCR purification kit (QIAGEN, Hilden, Germany). Libraries were prepared for sequencing using Nextera DNA

Library Preparation Kit (Illumina, FC-121). The libraries were purified using MinElute PCR purification kit (QIAGEN) and quantified using KAPA Library Quantification Kit (KAPA Biosystems). The libraries were multiplexed with a maximum of 24 samples per sequencing reaction.

**RNA-seq libraries**

BMDMs were lysed with TRIzol reagent (Life Technologies), and total RNA was purified using DIRECTzol RNA miniprep kit (Zymo Research). Strand-specific libraries were generated from 500 ng total RNA using KAPA Stranded mRNA-seq Library Preparation kit (KAPA Biosystems). Final libraries were checked for quality by agarose gel, quantified with Qubit, and multiplexed with a maximum of 24 samples per sequencing reaction.

**Sequencing and processing**

ChIP, ATAC, and RNA-seq libraries were single-end sequenced with a length of 50bp on an Illumina HiSeq 2500 at the UCLA Broad Stem Cell Research Center. The low quality 3'ends of reads were trimmed (cutoff q=30), and remaining adapter sequences were removed using cutadapt[6]. For ChIP and ATAC-seq, reads were aligned to the mm10 genome build using bowtie2[7] with default parameters except --very-sensitive and --non-deterministic options. For RNA-seq, reads were aligned to the mm10 genome using STAR[8] with the following options: --outFilterMultimapNmax 20, --alignSJoverhangMin 8, --alignSJDBoverhangMin 1, --outFilterMismatchNmax 999, --outFilterMismatchNoverLmax 0.04, --alignIntronMin 20, and --alignIntronMax 1000000 --seedSearchStartLmax 30. Aligned reads were filtered based on mapping score (MAPQ ≥ 30) by Samtools. For ChIP and ATAC-seq, duplicated reads were removed using Picard MarkDuplicates. Genome browser tracks for ChIP and ATAC-seq were generated using the bamCoverage function in deepTools[9] with the following options: --binSize 10, --smoothLength 30, --normalizeUsing RPGC, and for ChIP-seq only, --extendReads to average fragment length.

**ChIP-seq analysis**

MACS2 version 2.1.0[10] was used in broad mode to identify peaks for each sample using pooled input samples as control, FDR < 0.01, and extension size of average fragment length. These peaks were merged to generate a single reference peak file, and the number of reads that fell into each peak was counted using deeptools multiBamSummary[11] with extension size of average fragment length. edgeR[12] was used to perform normalization using the trimmed mean of M values (TMM) method and to construct a negative binomial model. For stimulus-specific dataset (Fig. 1), inducible peaks were identified in any stimulation condition compared to unstimulated by FDR < 0.05 and log2FC > 1. For IκB$\alpha$ KO vs control dataset (Fig. 4), inducible peaks were identified in TNF-treated samples compared to unstimulated by FDR < 0.05. Differentially inducible peaks between IκB$\alpha$ and KO were identified by FDR < 0.05 after re-constructing the negative binomial model to include only inducible peaks. Heat maps were generated with the pheatmap R package.

Analysis of transcription factor motif enrichment was performed using findMotifsGenome function in the HOMER suite[13], using the entire width of differential peaks as foreground and all detected peaks as background. Peaks of interest were annotated using HOMER annotatePeaks function with default parameters. ChipPeakAnno[14] was used to assess statistical significance of the overlap of differential peaks in the IκB$\alpha$ KO dataset and the stimulus-specific WT dataset.

RelA ChIP-seq data in Lipid A-stimulated BMDMs was previously published[15]. BAM files were obtained from Gene Expression Omnibus (accession number GSE67343). Peaks were called on each sample against input with FDR threshold < 0.01. A merged peak file was obtained, and overlaps between RelA ChIP-seq peaks and H3K4me1 ChIP-seq or ATAC-seq regions of interest were determined using the intersect function in the Bedtools suite[16].

**ATAC-seq analysis**

BAM files were analyzed with the R package csaw[17], using windows 200-nt wide, with 100-nt spacing. Windows with zero or background signal were removed. Sex-associated regions on chromosome X were also discarded[18]. Juxtaposed windows were merged in a single peak call up to a maximal chaining distance of 1,200 bp. Libraries were normalized using the TMM method. Batch effects were removed from the normalized count matrix with the RemoveBatchEffect function from the limma R package. Differential peaks were required to pass log2FC and FDR thresholds using csaw's Genewise negative binomial generalized linear models with quasi-likelihood tests, with batch effects controlled by modeling as blocking factors. Thresholds were 1 log2FC and 0.05 FDR for TNF-inducible peaks; and 0.5 log2FC and 0.1 FDR for IκB$\alpha$ KO vs control. For each peak call, the single 200-nt window with the highest abundance was retained as representative of that peak. Clustering was performed using the partitioning around medoids (PAM) algorithm on peak counts per million z-scaled across samples. Heat maps of peak signals were generated using the pheatmap R package. H3K4me1 signal from windows matching those selected from ATAC-seq analysis was extracted using csaw by extending each 200-nt ATAC-seq window by additional 200 nt from 5' end and 200 nt from 3'end, normalized using TMM offsets computed genome-wide.

To obtain a normalized percentage of cells with open chromatin (Fig. 3g), ATAC counts were normalized as follows: over the 322 regions of interest, the 0.1 percentile of ATAC-seq signal was assigned a value of "0% of cells accessible." The 99.9 percentile of ATAC-seq signal was assigned a value of "100% of cells accessible," and all data points were scaled to these values.

**RNA-seq analysis**

Transcript read counts were computed by the featureCounts function in the Subread package[19]. TMM normalization was performed using the edgeR package. Genes below an expression threshold of 4.6 CPM in all samples were excluded from downstream analysis. Inducible genes were identified in TNF-treated samples compared to unstimulated by FDR < 0.05. Differentially inducible genes between IκB$\alpha$ and KO were identified by FDR < 0.05 after re-constructing the negative binomial model to include only inducible genes. Heat maps were generated with the pheatmap R package. Gene ontology analysis was performed using the PANTHER database[20] with all expressed genes as background

**Model of nucleosome opening**

*Model formulation:* A multistep model of DNA unwrapping from the histone octamer was formulated based on structural studies that identified 14 contacts between the histone octamer and double helical DNA and biophysical studies of single nucleosomes *in vitro* that revealed multiple, step-wise transitions in DNA unwrapping[21–23]. The model describes the population average or probability of many stochastic events, with each species representing the fraction of NFκB-responsive latent enhancers in a cell (*E*) at a given state of accessibility. $E_{-14}$ describes the most closed state in which all 14 contact points are engaged, and $E_0$ the most open state in which the histone octamer is entirely evicted.

$$E_{-14} \underset{\alpha_{-14}}{\overset{\beta_{-14}(t)}{\rightleftharpoons}} E_{-13} \cdots \underset{\alpha_{i}}{\overset{\beta_{i}(t)}{\rightleftharpoons}} E_{i+1} \cdots \underset{\alpha_{-1}}{\overset{\beta_{-1}(t)}{\rightleftharpoons}} E_{0}$$

A system of ordinary differential equations was formulated to describe this model:

$$\frac{dE_{-14}}{dt} = \alpha_{-14}E_{-13} - \beta_{-14}(t)E_{-14} \qquad (1)$$

$$\frac{dE_{-13}}{dt} = \beta_{-14}(t)E_{-14} - \alpha_{-14}E_{-13} - \beta_{-13}(t)E_{-13} + \alpha_{-13}E_{-12} \qquad (2)$$

...

$$\frac{dE_{i}}{dt} = \beta_{i-1}(t)E_{i-1} - \alpha_{i-1}E_{i} - \beta_{i}(t)E_{i} + \alpha_{i}E_{i+1} \qquad (3)$$

...

$$\frac{dE_{0}}{dt} = \beta_{-1}(t)E_{-1} - \alpha_{-1}E_{0} \qquad (4)$$

The rate constant $\alpha_i$ describes the closing transition. The term $\beta_i(t)$ represents an NFκB-dependent opening transition that varies with time as nuclear NFκB concentration varies with time. Experimental studies have demonstrated that NFκB can interact with nucleosomal DNA[24] while the lowest energy structure of the NFκB-DNA complex is sterically incompatible with DNA-histone octamer interactions[25,26], suggesting that NFκB can promote nucleosome unwrapping. The opening transition is thus formulated as:

$$\beta_i(t) = \frac{k_i[NF\kappa B(t)]^n}{K_D^n + [NF\kappa B(t)]^n}$$

where $K_D$ is the dissociation constant of the NFκB -DNA interaction, $n$ is the Hill coefficient, $k_i$ is the transition rate constant, and [NFκB*(t)*] represents the time-dependent nuclear NFκB concentration.

*Model parameters:* $K_D$ and $n$ were surveyed across reasonable ranges, producing qualitatively similar results (Extended Data Fig. 4a,b). For the shown simulations $K_D$ was set to 0.025 $\mu M$ and the Hill coefficient was set to three. The rate of nucleosome unwrapping was also surveyed across a wide range (Extended Data Fig. 4c), and the ratio of unwrapping to rewrapping rates was selected based on biophysical measurements[23]. Based on this literature, the initial rewrapping rate is approximately 5-10 times faster than the unwrapping rate, so we set the ratio of rewrapping to unwrapping to 7.5 at the first step. As *in vivo* nucleosomes are stabilized by linker histones and cooperative binding within the nucleosome array, we scaled transition rates to be 50-fold slower than *in vitro* measurements, with the unwrapping rate constant of the first opening step $k_{-14}$ set to 10 $\mu M^{-1} min^{-1}$, and the rewrapping rate constant $\square_{-14}$ to 75 $min^{-1}$. To account for the inherent cooperativity of contact points within a nucleosome[27], stepwise increases in unwrapping rates and decreases in rewrapping rates were included. These cooperativity factors were examined through a parameter sweep (Extended Data Fig. 4d,e), and cooperativity factors of 1.2 for unwrapping and 0.8 for rewrapping were selected.

*Model simulations:* Simulations were performed in MATLAB R2014b. Experimental and theoretical single cell traces (Supplemental Table 1) were used as input to the ODE system described above. Experimental values of mVenus-RelA fluorescent intensities were converted to $\mu M$ concentrations of NFκB based on previously published NFκB models, with the maximum fluorescence of the first peak for the single cell trajectories approximating the maximal nuclear NFκB concentration of 0.25 μM reported in previous studies[28]. For simulations, the initial value of the $E_{-14}$ state was set to 1, and all other states set to 0. The MATLAB function *ode*15*s* was used to solve the ODE system, and the concentration of the most open chromatin state $E_0$ was plotted.

***References***


1. Taylor B, Adelaja A, Hoffmann A. Deciphering the language of immune sentinel cells to signal inflammatory threats. BioRxiv. 2020;

2. Shih VF-S, Kearns JD, Basak S, Savinova OV, Ghosh G, Hoffmann A. Kinetic control of negative feedback regulators of NF- B/RelA determines their pathogen- and cytokine-receptor signaling specificity. Proceedings of the National Academy of Sciences. 2009 Jun 16;106(24):9619–9624.

3. Kearns JD, Basak S, Werner SL, Huang CS, Hoffmann A. IkappaBepsilon provides negative feedback to control NF-kappaB oscillations, signaling dynamics, and inflammatory gene expression. J Cell Biol. 2006 Jun 5;173(5):659–664. PMCID: PMC2063883

4. Peng B, Ling J, Lee AJ, Wang Z, Chang Z, Jin W, Kang Y, Zhang R, Shim D, Wang H, Fleming JB, Zheng H, Sun S-C, Chiao PJ. Defective feedback regulation of NF-kappaB underlies Sjogren's syndrome in mice with mutated kappaB enhancers of the IkappaBalpha promoter. Proc Natl Acad Sci USA. 2010 Aug 24;107(34):15193–15198. PMCID: PMC2930541

5. Selimkhanov J, Taylor B, Yao J, Pilko A, Albeck J, Hoffmann A, Tsimring L, Wollman R. Accurate information transmission through dynamic biochemical signaling networks. Science. 2014 Dec 12;346(6215):1370–1373.

6. Martin M. Cutadapt removes adapter sequences from high-throughput sequencing reads. EMBnet j. 2011 May 2;17(1):10.

7. Langmead B, Salzberg SL. Fast gapped-read alignment with Bowtie 2. Nature Methods. 2012 Mar 4;9(4):357–9. PMCID: PMC3322381

8. Dobin A, Davis CA, Schlesinger F, Drenkow J, Zaleski C, Jha S, Batut P, Chaisson M, Gingeras TR. STAR: ultrafast universal RNA-seq aligner. Bioinformatics. 2013;29(1):15–21.

9. Ramírez F, Ryan DP, Grüning B, Bhardwaj V, Kilpert F, Richter AS, Heyne S, Dündar F, Manke T. deepTools2: a next generation web server for deep-sequencing data analysis. Nucleic Acids Res. 2016 Jul 8;44(W1):W160–W165.

10. Zhang Y, Liu T, Meyer CA, Eeckhoute J, Johnson DS, Bernstein BE, Nusbaum C, Myers RM, Brown M, Li W, Liu XS. Model-based analysis of ChIP-Seq (MACS). Genome biology. 2008;9(9):R137. PMCID: PMC2592715

11. Ramírez F, Ryan DP, Grüning B, Bhardwaj V, Kilpert F, Richter AS, Heyne S, Dündar F, Manke T. deepTools2: a next generation web server for deep-sequencing data analysis. Nucleic Acids Res. 2016 Jul 8;44(W1):W160–W165.

12. Robinson MD, McCarthy DJ, Smyth GK. edgeR: a Bioconductor package for differential expression analysis of digital gene expression data. Bioinformatics. 2010 Jan 1;26(1):139–40. PMCID: PMC2796818

13. Heinz S, Benner C, Spann N, Bertolino E, Lin YC, Laslo P, Cheng JX, Murre C, Singh H, Glass CK. Simple combinations of lineage-determining transcription factors prime cis-regulatory elements required for macrophage and B cell identities. Mol Cell. 2010 May 28;38(4):576–589. PMCID: PMC2898526

14. Zhu LJ, Gazin C, Lawson ND, Pagès H, Lin SM, Lapointe DS, Green MR. ChIPpeakAnno: a Bioconductor package to annotate ChIP-seq and ChIP-chip data. BMC Bioinformatics. 2010 Dec;11(1):237.

15. Tong A-J, Liu X, Thomas BJ, Lissner MM, Baker MR, Senagolage MD, Allred AL, Barish GD, Smale ST. A Stringent Systems Approach Uncovers Gene-Specific Mechanisms Regulating Inflammation. Cell. 2016 Mar 24;165(1):165–179. PMCID: PMC4808443

16. Quinlan AR, Hall IM. BEDTools: a flexible suite of utilities for comparing genomic features. Bioinformatics. 2010 Mar 15;26(6):841–2. PMCID: PMC2832824

17. Lun ATL, Smyth GK. csaw: a Bioconductor package for differential binding analysis of ChIP-seq data using sliding windows. Nucleic Acids Res. 2016 Mar 18;44(5):e45. PMCID: PMC4797262

18. Mognol GP, Spreafico R, Wong V, Scott-Browne JP, Togher S, Hoffmann A, Hogan PG, Rao A, Trifari S. Exhaustion-associated regulatory regions in CD8 $^{+}$ tumor-infiltrating T cells. Proc Natl Acad Sci USA. 2017 Mar 28;114(13):E2776–E2785.

19. Liao Y, Smyth GK, Shi W. featureCounts: an efficient general purpose program for assigning sequence reads to genomic features. Bioinformatics. 2014 Apr 1;30(7):923–930. PMID: 24227677

20. Mi H, Muruganujan A, Ebert D, Huang X, Thomas PD. PANTHER version 14: more genomes, a new PANTHER GO-slim and improvements in enrichment analysis tools. Nucleic Acids Res. 2019 Jan 8;47(D1):D419–D426. PMCID: PMC6323939

21. Möbius W, Neher RA, Gerland U. Kinetic Accessibility of Buried DNA Sites in Nucleosomes. Phys Rev Lett. 2006 Nov 14;97(20):208102.

22. Luger K, Mäder AW, Richmond RK, Sargent DF, Richmond TJ. Crystal structure of the nucleosome core particle at 2.8 A resolution. Nature. 1997 Sep 18;389(6648):251–260. PMID: 9305837

23. Tims HS, Gurunathan K, Levitus M, Widom J. Dynamics of Nucleosome Invasion by DNA Binding Proteins. Journal of Molecular Biology. 2011 Aug;411(2):430–448.

24. Lone IN, Shukla MS, Charles Richard JL, Peshev ZY, Dimitrov S, Angelov D. Binding of NF-κB to Nucleosomes: Effect of Translational Positioning, Nucleosome Remodeling and Linker Histone H1. Schübeler D, editor. PLoS Genet. 2013 Sep 26;9(9):e1003830.

25. Chen FE, Huang DB, Chen YQ, Ghosh G. Crystal structure of p50/p65 heterodimer of transcription factor NF-kappaB bound to DNA. Nature. 1998 Jan 22;391(6665):410–413. PMID: 9450761

26. Suto RK, Clarkson MJ, Tremethick DJ, Luger K. Crystal structure of a nucleosome core particle containing the variant histone H2A.Z. Nat Struct Biol. 2000 Dec;7(12):1121–1124. PMID: 11101893

27. Miller JA, Widom J. Collaborative competition mechanism for gene activation in vivo. Mol Cell Biol. 2003 Mar;23(5):1623–1632. PMCID: PMC151720

28. Shih VF-S, Davis-Turak J, Macal M, Huang JQ, Ponomarenko J, Kearns JD, Yu T, Fagerlund R, Asagiri M, Zuniga EI, Hoffmann A. Control of RelB during dendritic cell activation integrates canonical and noncanonical NF-κB pathways. Nat Immunol. 2012 Dec;13(12):1162–1170.
