## Extended Data Figures for "NFκB dynamics determine the stimulus-specificity of epigenomic reprogramming in macrophages"

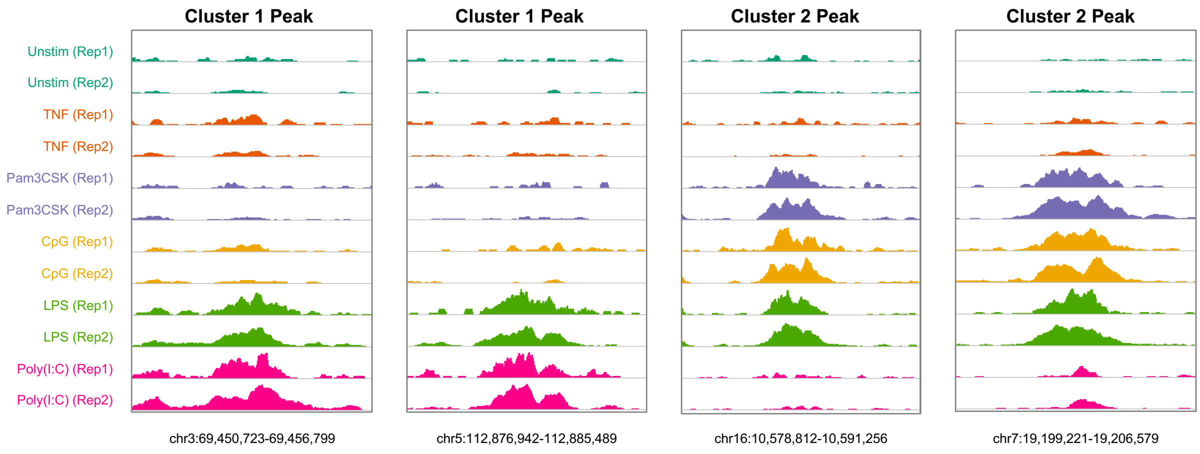


**Extended data, Figure 1:** Genome browser tracks of representative stimulus-specific *de novo* enhancers from Fig. 1a., two replicates per condition.

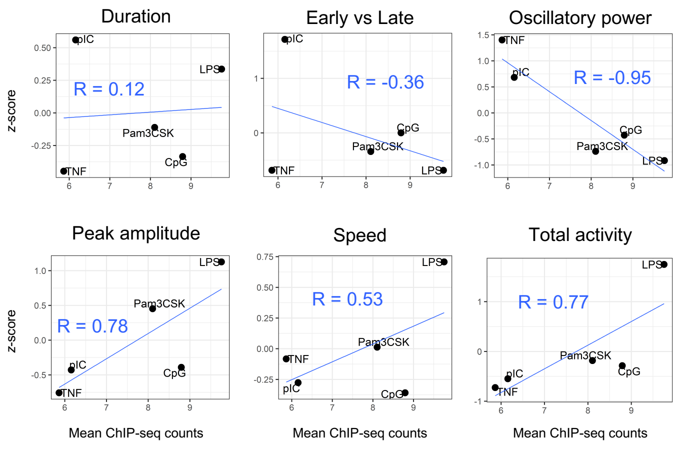


**Extended data, Figure 2: Correlation of NFκB dynamics to ChIP-seq data.** Scatterplots and correlations of mean Cluster 2 ChIP-seq counts *vs.* stimulus-specific z-scores for each of the six key features of NFκB signaling dynamics

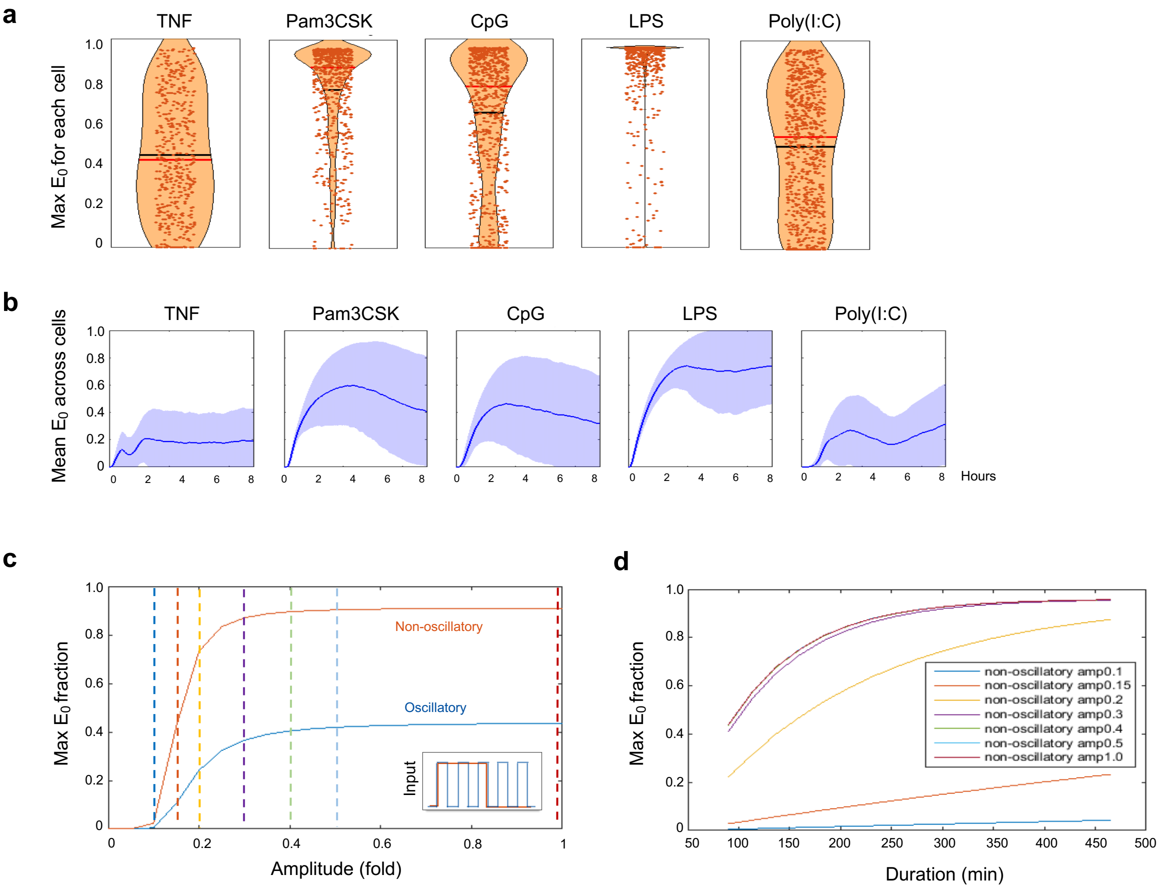


**Extended data, Figure 3: Supplemental model simulations. a)** Violin plots of maximum chromatin opening over eight hours per single-cell stimulation, using NFκB trajectories as input to the model. Black line = mean, Red line = median. **b)** Simulated mean chromatin opening over time across all single cells. **c)** Model simulations across a range of NFκB amplitudes, comparing oscillatory and non-oscillatory trajectories. **d)** Model simulations across a range of NFκB durations, comparing a range of NFκB amplitudes marked by dotted lines in (c).

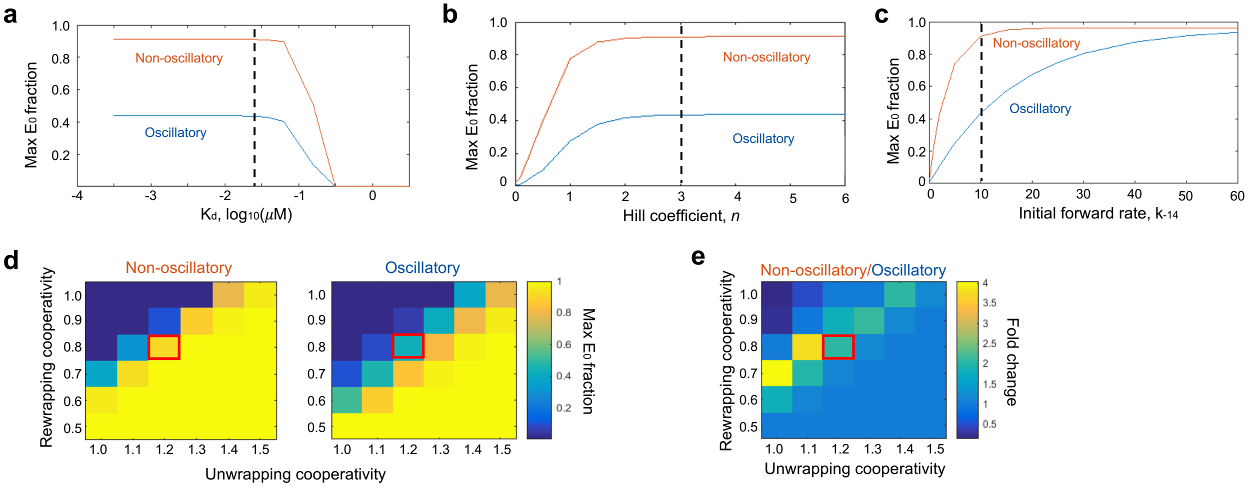


**Extended data, Figure 4: Parameter sensitivity analysis. (a)** Chromatin opening behavior when the model is tested across a range of $K_D$s, **(b)** across a range of Hill coefficients, or **(c)** across a range of forward rates for the first unwrapping step, $k_{-14}$. For model simulations (Fig. 2d), $K_D$ = 0.025, Hill = 3, and $k_{-14}$ = 10 were used, marked by the dotted black line. **(d-e)** Heat map of chromatin opening across a range of unwrapping and rewrapping cooperativity factors, showing maximum $E_0$ fraction in non-oscillatory and oscillatory conditions (d) and fold change difference between maximum non-oscillatory and oscillatory conditions (e). Red box indicates the parameter values used for model simulations.

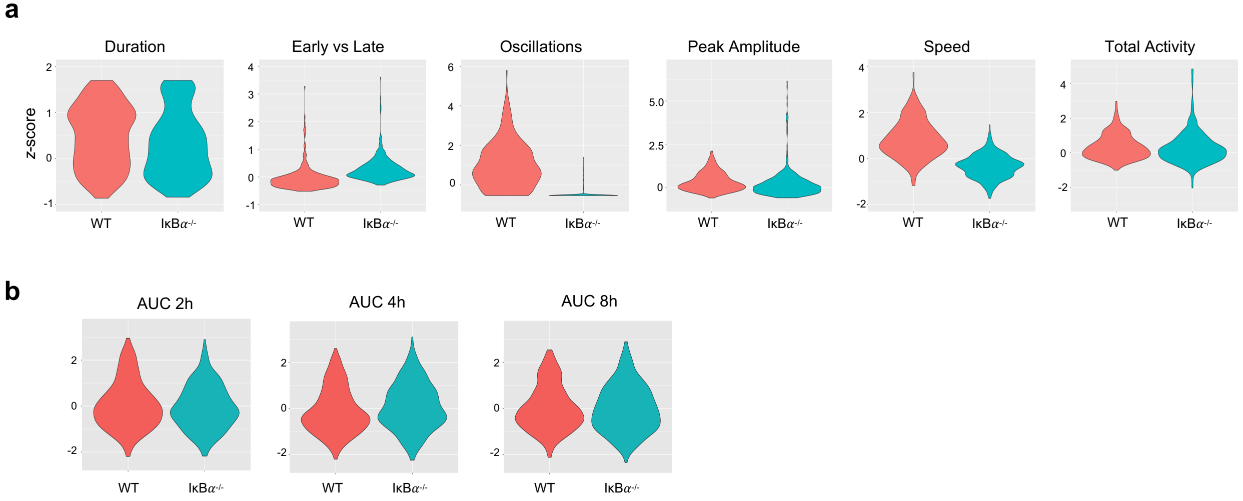


**Extended data, Figure 5: NFκB dynamics in TNF-stimulated IκB$\alpha^{-/-}$ vs WT BMDMs. (a)** Violin plots of single-cell distributions for the six key NFκB signaling features. **(b)** Violin plots of single-cell distributions for areas under the NFκB activity curve at two, four, and eight hours. **(c)** Bar graph of Kolmogorov–Smirnov (K-S) test statistic for difference between distributions of IκB$\alpha^{-/-}$ and WT cells for key signaling features and areas under the curve. Directionality of bars is based on whether observed differences would intuitively favor greater NFκB activity in WT or knockout cells.

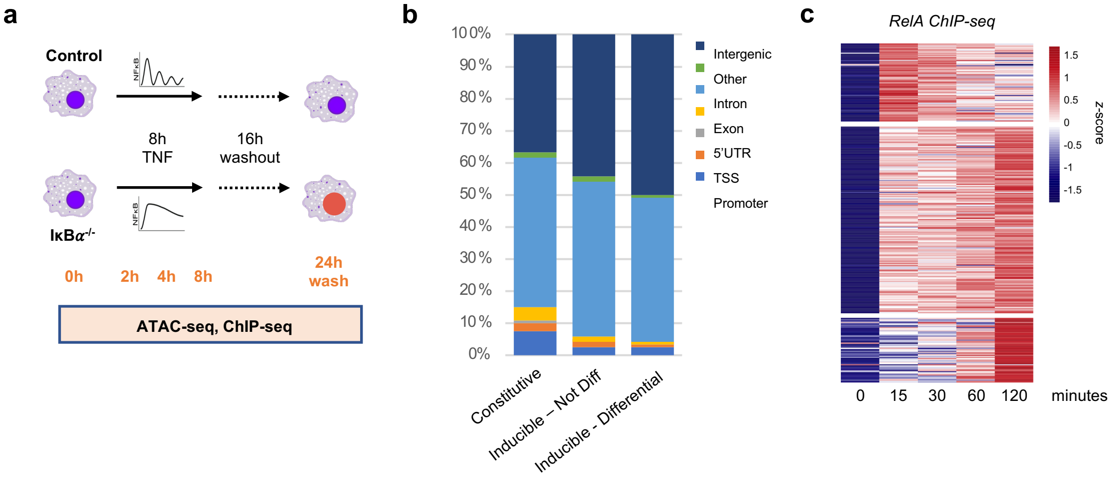


**Extended data, Figure 6: Supplemental ATAC-seq data. a)** Schematic of ATAC and ChIP-seq experiments in IκB$\alpha^{-/-}$ and control BMDMs. **b)** Genomic distribution of three categories of accessible regions identified by ATAC-seq. **c)** Heat map of Lipid A-stimulated NFκB RelA ChIP-seq signal[25] at 322 inducible-differential ATAC-seq regions, 311 of which overlap with a RelA ChIP-seq peak.

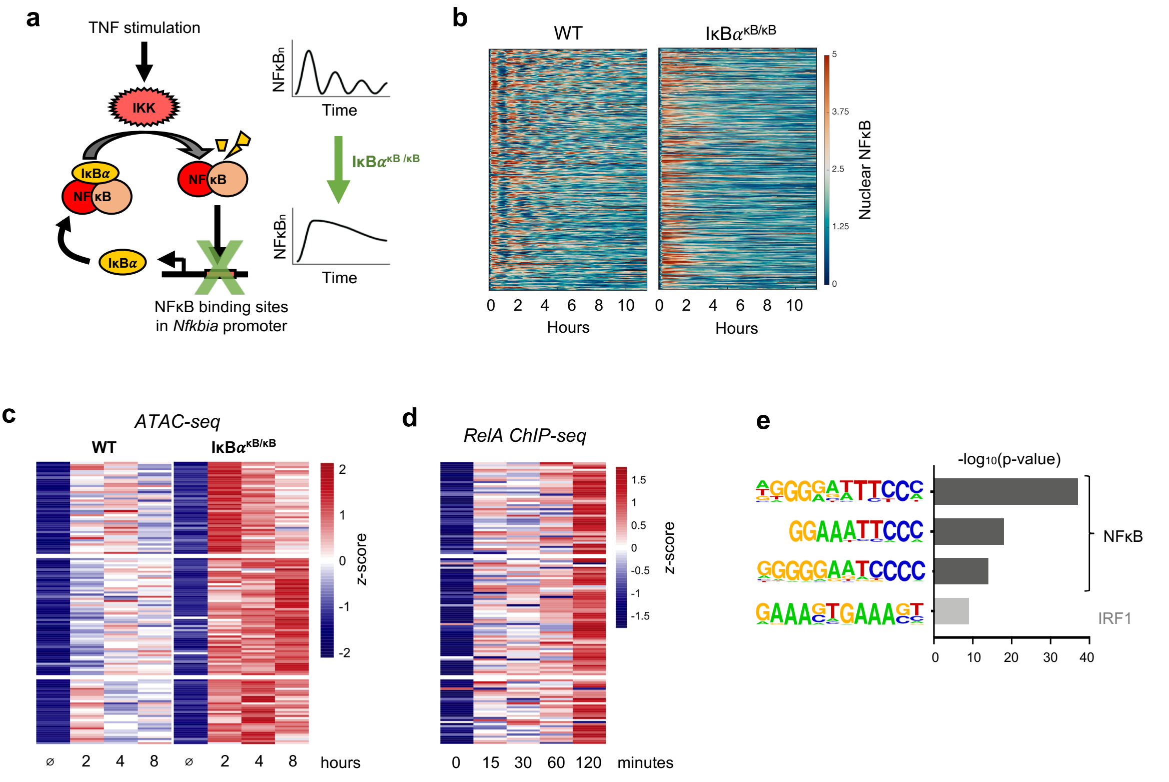


**Extended data, Figure 7: *Nfkbia*$^{\kappa B/\kappa B}$ mutant as a complementary model of non-oscillatory NFκB. a)** Schematic of *Nfkbia*$^{\kappa B/\kappa B}$ mutation, abolishing inducible IκB*α* by disrupting NFκB binding sites in promoter[26]. **b)** Heat map of single cell NFκB trajectories by microscopy, comparing TNF response in WT *vs.* IκB*α*$^{\kappa B/\kappa B}$ BMDMs. **c)** Heat map of ATAC-seq signal at 131 genomic regions that are TNF-inducible and differential between IκB*α*$^{\kappa B/\kappa B}$ and WT. **d)** Heat map of Lipid-A stimulated NFκB RelA ChIP-seq signal[25] at 131 inducible-differential ATAC-seq regions, 118 of which overlap with a RelA ChIP-seq peak. **e)** Known transcription factor motifs with greatest enrichment in differentially inducible ATAC-seq regions.

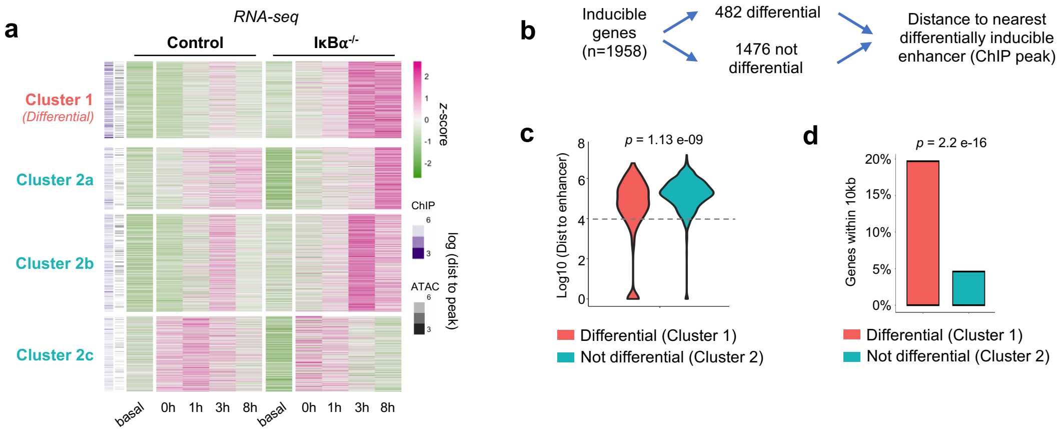


**Extended data, Figure 8: Gene-centric approach to investigate the function of dynamics-dependent enhancers. a)** Heat map of TNF-inducible genes in control or IκB$\alpha^{-/-}$ BMDMs (n=1958). Cluster 1 genes are differentially induced between IκB$\alpha^{-/-}$ and control. Clusters 2a-2c genes are inducible but not differential. **b)** Gene-centric approach to map inducible genes to nearest dynamics-dependent enhancers. **c)** Violin plots of distance from gene TSS to nearest differentially inducible enhancer, comparing genes in Cluster 1 (differential) and Cluster 2 (not differential) by K-S test. **d)** Percentage of genes in each cluster within 10kb (dashed line, panel c) of a differentially inducible enhancer, evaluated by *chi* square test.
